# *DMC1* stabilizes synapsis and crossover at high and low temperatures during wheat meiosis

**DOI:** 10.1101/2023.04.18.537302

**Authors:** Tracie N. Draeger, María-Dolores Rey, Sadiye Hayta, Mark Smedley, Azahara C. Martín, Graham Moore

## Abstract

Effective chromosome synapsis and crossover during meiosis are essential for fertility, especially in grain crops such as wheat. These processes function most efficiently in wheat at temperatures between 17-23 °C, although the genetic mechanisms for such temperature dependence are unknown. In a previously identified mutant of the hexaploid wheat reference variety ‘Chinese Spring’ lacking the long arm of chromosome 5D, exposure to low temperatures during meiosis resulted in asynapsis and crossover failure. In a second mutant (*ttmei1*), containing a 4 Mb deletion in chromosome 5DL, exposure to 13 °C led to similarly high levels of asynapsis and univalence. Moreover, exposure to 30 °C led to a significant, but less extreme effect on crossover. Previously, we proposed that, of 41 genes deleted in this 4 Mb region, the major meiotic gene *TaDMC1-D1* was the most likely candidate for preservation of synapsis and crossover at low (and possibly high) temperatures. In the current study, using RNA-guided Cas9, we developed a new Chinese Spring CRISPR mutant, containing a 39 bp deletion in the 5D copy of *DMC1*, representing the first reported CRISPR-Cas9 targeted mutagenesis in Chinese Spring, and the first CRISPR mutant for *DMC1* in wheat. In controlled environment experiments, wild-type Chinese Spring, CRISPR *dmc1-D1* and backcrossed *ttmei1* mutants were exposed to either high or low temperatures during the temperature-sensitive period from premeiotic interphase to early meiosis I. After 6-7 days at 13 °C, crossover decreased by over 95% in the *dmc1-D1* mutants, when compared with wild-type plants grown under the same conditions. After 24 hours at 30 °C, *dmc1-D1* mutants exhibited a reduced number of crossovers and increased univalence, although these differences were less marked than at 13 °C. Similar results were obtained for *ttmei1* mutants, although their scores were more variable, possibly reflecting higher levels of background mutation. These experiments confirm our previous hypothesis that *DMC1-D1* is responsible for preservation of normal synapsis and crossover at low and, to a certain extent, high temperatures. Given that reductions in crossover have significant effects on grain yield, these results have important implications for wheat breeding, particularly in the face of climate change.

**Key message:** The meiotic recombination gene *DMC1* on wheat chromosome 5D preserves normal chromosome synapsis and crossover during periods of high and low temperature.

## Introduction

Like many plants, wheat (*Triticum aestivum* L) is highly sensitive to temperatures that fall outside the range typically experienced during a growing season. The optimum temperature range for wheat growth over an entire season is generally considered to be around 17-23 °C, with temperatures above or below this range significantly reducing grain yield (Porter and Gawith, 1999). The extent to which temperature stresses affect yield is dependent on developmental stage, with reproductive stages more sensitive to high temperatures than vegetative growth stages (Fisher and Maurer, 1976). Even quite short periods of high temperature (20-24 hours at 30 °C) during meiosis can reduce grain number (Saini and Aspinall, 1982; Draeger and Moore, 2017). Meiosis has also been identified as the stage most sensitive to low temperature stress (Thakur et al., 2010), with reductions in grain yield occurring when low temperatures coincide with the ‘booting’ stage, which broadly corresponds to meiosis (Ji et al., 2017).

Meiosis is a highly dynamic process during which parental chromosomes pair and recombine. It is essential for gamete formation in sexually reproducing organisms. During early meiosis, the parental chromosomes align alongside each other as pairs, enabling a proteinaceous structure, the synaptonemal complex (SC), to assemble between them, linking the chromosomes along their entire lengths in a process called synapsis (Page and Hawley, 2004). The SC is thought to provide the structural framework for meiotic recombination to take place. Recombination initiates by programmed double-strand breaks (DSBs) in the DNA. A small minority of these are repaired as crossovers, forming physical connections between each chromosome pair, enabling genetic information to be reciprocally exchanged, and potentially creating new advantageous allelic combinations. These physical connections can be seen cytologically as chiasmata. Bread wheat is an allohexaploid (2n = 6x = 42), made up of three related (homeologous) sub-genomes (A, B and D). During wheat meiosis, chromosome behaviour is tightly controlled by the *TaZIP4-B2 (Ph1)* gene, which promotes synapsis and crossover between homologous chromosomes (homologs), rather than between homeologous chromosomes (homeologs) from related genomes (Riley and Chapman, 1958; Sears and Okamoto, 1958; Martín et al., 2014 and 2017; Rey et al., 2017; Draeger et al., 2023). This allows the three genomes to behave as separate diploids during meiosis, thus maintaining genome stability and fertility.

Assembly of the SC completes at pachytene, and by diplotene crossovers are fully formed. The SC then disassembles, leaving homologs connected only by their chiasmata. At metaphase I, the homolog pairs align on the equatorial plate, and the chiasmata connecting the chromosomes can be seen using light microscopy. At this stage, in hexaploid wheat, the chromosomes of each parent are normally bound together as 21 bivalent pairs, most forming as ring bivalents with two chiasmata linking them, one in each arm, usually towards the distal ends. Occasionally, the homologs are bound together by chiasmata in one arm only, forming a rod bivalent. In wheat, on average, there are around 2.3 crossovers per homolog pair (Miller and Reader, 1985). At least one crossover (the ‘obligate’ crossover) must form between each bivalent pair to ensure accurate chromosome segregation and balanced gametes in daughter cells (Zickler and Kleckner, 1999; Jones and Franklin, 2006), which is vital for maintaining genome stability and fertility.

Assembly of the SC is a highly temperature-sensitive process (Bilgir et al., 2013). Both high and low temperatures can lead to disruption of synapsis, resulting in unpaired univalent chromosomes that segregate randomly or are lost completely (reviewed in Bomblies et al., 2015 and Morgan et al., 2017). Relatively small changes in temperature can alter the frequency and distribution of crossovers (Elliott, 1955; Dowrick, 1957; Bayliss and Riley, 1972a; Higgins et al., 2012). In wheat, high and low temperatures have been found to generally decrease chiasma frequency. In the wheat cultivar ‘Chinese Spring’, this temperature sensitivity is under the genetic control of a major gene located on chromosome 5D (Riley, 1966). In Chinese Spring plants lacking chromosome 5D, numbers of chiasmata progressively decrease as the temperature falls below the optimum range (Riley, 1966; Bayliss and Riley, 1972a). Chiasma frequency is greatly reduced at 15 °C, and pronounced chromosome pairing failure occurs at 12 °C, resulting in complete male sterility (Riley, 1966; Hayter and Riley, 1967). This reduction in chiasma frequency is due to failure of chromosome pairing at zygotene, although the temperature-sensitive phase is earlier, during premeiotic interphase, prior to DNA synthesis (Bayliss and Riley, 1972b).

These studies led to the proposal that there must be a gene on chromosome 5D that stabilizes chromosome pairing at low temperatures. This putative gene, named *low-temperature pairing* (*Ltp*) (Hayter and Riley,1967), was further defined to the long arm of chromosome 5D (Hayter, 1969) and later renamed *Ltp1* (Queiroz et al., 1991). In plants lacking chromosome 5D, chiasma frequency also appears to decrease progressively at temperatures of ∼30 °C and above (Bayliss and Riley, 1972a), suggesting that chromosome 5D may also be associated with tolerance to high temperatures. However, this suggestion was based on the scoring of a few cells only, because high temperature treatments for 3 days made the chromosomes too sticky for accurate scoring.

More recently, we used wheat (Chinese Spring) lines with terminal deletions of 5DL to delimit the *Ltp1* locus to the proximal half of chromosome 5DL (Draeger et al., 2020). KASP markers specific to 5DL, that mapped within this delimited region, were used to screen ∼2500 gamma-irradiated deletion lines, from which 16 plants were identified with 5DL deletions. Mapping and candidate gene identification were facilitated by resources including the Chinese Spring IWGSC RefSeq v1.0 genome assembly (International Wheat Genome Sequencing Consortium, 2018), the Wheat 820 K Axiom^®^ Breeders’ Array probe set (Winfield et al., 2015) and the Ensembl Plants database (Bolser et al., 2016). The 16 mutant plants were then exposed to low temperature (13 °C) for 7 days, during a period lasting from premeiotic interphase to early meiosis I. From this, we identified a deletion mutant with meiotic chromosomes exhibiting extremely high levels of asynapsis and chromosome univalence after the low temperature treatment. This was very similar to the phenotype previously described for *Ltp1* when the whole of chromosome 5D was absent. Exposure of this same *ltp1-like* deletion mutant to 30 °C for 24 hours during the same developmental period, also led to a reduced number of crossovers and increased univalence, although the effect was less pronounced than that observed following exposure to 13 °C. However, as the deletion had a clear effect on chromosome pairing at 30 °C as well as at 13 °C, we renamed the mutant line *temperature tolerant meiosis 1 (ttmei1*), to reflect its reduced tolerance to high temperature in addition to its loss of the low temperature pairing gene *Ltp1*.

Using KASP genotyping, we then mapped the *ttmei1* deletion to a 4 Mb region of chromosome 5DL. Of the 41 genes deleted in this region, 18 were expressed during meiosis, of which 12 were high confidence genes. Of these, the strongest candidate for the observed effects on meiosis was the D-genome homeolog of the meiotic recombination gene *DMC1* (*TaDMC1-D1*), which had a ten-fold higher expression level in meiotic tissues compared with non-meiotic. For the remaining 17 genes expressed during meiosis, expression was proportionally higher in non-meiotic tissues, and none had any previously known meiotic function. Moreover, *TaDMC1-D1* was expressed most highly during early prophase I, which in wheat coincides with the ‘telomere bouquet’ stage, when synapsis is initiated (Martín et al., 2017). Consistent with this, in the *ttmei1* mutant, synapsis was abnormal after exposure to 13 °C and did not complete. Therefore, we proposed that *DMC1-D1* was probably responsible for the *Ltp1/ttmei1* phenotype.

DMC1 (Disrupted Meiotic cDNA 1) is a recombinase that plays a central role in meiotic recombination, performing homology search, strand invasion and strand exchange during repair of meiotic DSBs. The process of strand invasion is fundamentally dynamic, which makes it vulnerable to disruption by temperature stress. In the current study, we have developed a CRISPR *dmc1-D1* mutant in the hexaploid wheat variety Chinese Spring. Until recently, very few wheat genotypes have been transformable, but the development of a GRF-GIF chimeric protein (Debernadi et al., 2020) has addressed this issue, and this technology, combined with our efficient transformation system (Hayta et al., 2019, 2021), has allowed us to generate the first CRISPR mutants in Chinese Spring. Chinese Spring is used as the hexaploid wheat reference genome by most wheat researchers, as it has a fully annotated sequenced genome (International Wheat Genome Sequencing Consortium, 2018), with all data integrated into the Ensembl Plants database (Bolser et al., 2016).

We have also backcrossed *ttmei1* mutants with wild-type Chinese Spring plants, to reduce background mutations. We have exposed the *ttmei1* and CRISPR *dmc1-D1* mutants to high (30 °C) and low (13 °C) temperatures during the sensitive period of premeiosis to meiosis I, to determine whether the D-genome copy of *DMC1* has a stabilizing effect on chromosome synapsis and crossover in wheat.

## Materials and Methods

### Plant materials

Mutants for *DMC1-D1* (TraesCS5D02G141200) and *TTMEI1* were derived from the hexaploid wheat cultivar ‘Chinese Spring’ (*Triticum aestivum L*., 2n = 6x = 42; AABBDD), which was also used as a wild-type control. The CRISPR *Tadmc1-D1* mutant was developed within the Crop Transformation Facility (BRACT) at the John Innes Centre, using RNA-guided Cas9. The *Tattmei1* deletion mutant was generated previously by gamma irradiation (Draeger et al., 2020). *Tattmei1* has a 4 Mb interstitial deletion on chromosome 5DL, with 41 genes deleted, including *TaDMC1-D1*. For the current study, *ttmei1* M_2_ plants were backcrossed twice with Chinese Spring to produce Bc_2_F_2_ plants with fewer background mutations.

### Production of *Tadmc1-D1* CRISPR mutants using RNA-guided Cas9 Plasmid assembly

The Chinese Spring genomic DNA sequences of *TaDMC1* homeologs were aligned using the software Geneious Prime, version 2020.2.4 (Biomatters). Two guide RNAs were designed targeting the D genome copy of *TaDMC1*, denoted as TaDMC1-D Guide1: 5’-GCTCATGGAGGCCGACCGGG-3’ and TaDMC1-D Guide2: 5’-CAAGCAGCTCATCAAGCGTT-3’. The selected guides were ordered as forward and complementary oligonucleotides, with 5’ overhangs to enable cloning. Using standard Golden Gate MoClo assembly (Werner et al., 2012), the guide RNAs were cloned between the TaU6 promoter and the guide scaffold for *Streptococcus pyogenes* Cas9, in the Level 1 acceptor plasmids pL1P3-TaU6 (Addgene #165599) and pL1P4-TaU6 (Addgene #165600), as described in Smedley et al., (2021). These Level 1 plasmids were sequenced, before proceeding to the Level 2 assembly. Level 2 assembly was performed using the Level 2 acceptor pGoldenGreenGate-M (pGGG-M) (Addgene #165422) binary vector (Smedley et al., 2021). The Level 1 plasmids pL1P1OsActinP:*hpt*-int:35sT selection cassette (Addgene #165423), pL1P2OsUbiP:Cas9:NosT (Addgene #165424), pL1P5ZmUbiP:GRF-GIF:NosT

(Addgene #198046) and guide cassettes were assembled into pGGG-M along with end linker pELE-5 (Addgene #48020). The resulting plasmid was named pGGG-TaDMC1-D and sequenced to ensure authenticity before transferring to *Agrobacterium*.

### Preparation of Agrobacterium for transformation

The hypervirulent *Agrobacterium tumefaciens* strain AGL1 (Lazo et al., 1991) was used for the wheat transformation experiments. The pGGG-TaDMC1-D vector was electroporated into *A. tumefaciens* AGL1 competent cells, as previously described (Hayta et al., 2021), and was co-electroporated with the helper plasmid pAL155 (Hayta et al., 2019). Single colonies of *Agrobacterium* AGL1, containing the pGGG-TaDMC1-D vector, were inoculated into 10 ml of LB (Bertani, 1951) liquid medium containing kanamycin (50 μg mL^-1^) and rifampicin (50 μg mL^-1^), and incubated at 28 °C, shaking at 200 rpm for ∼ 65 hours. A modified method of Tingay et al., (1997) was used to prepare *Agrobacterium* standard inoculums for transformation, as previously described by Hayta et al., (2021). Briefly, equal quantities of 30% sterile glycerol and *Agrobacterium* culture were mixed by inverting, producing 400 μl aliquots of standard inoculums in 0.5 ml Eppendorf tubes which were then frozen and stored at -80 °C until required.

### Agrobacterium transformation of Chinese Spring

Wheat transformation was performed as previously published by Hayta et al., (2021), with slight modification. The construct incorporated GRF4-GIF1 technology (Debernardi et al., 2020). Briefly, Chinese Spring plants were grown in a controlled environment room under a long-day photoperiod (16 h at 600 μmol m^−2^ s^−1^ light, at 20 °C day and 16 °C night). Wheat spikes were collected ∼14 days post anthesis (early milk stage GS73) when immature embryos were 1-1.5 mm in diameter. Under aseptic conditions, immature embryos were isolated from surface sterilized grain.

Isolated immature embryos were pre-treated by centrifugation in liquid medium, prior to *Agrobacterium* inoculation. Embryos were transferred to co-cultivation medium, scutellum side up, and incubated at 24 °C in the dark for 3 days co-cultivation. Embryogenic axes were excised and discarded, before the embryos were transferred to wheat callus induction (WCI) medium without selection, for 5 days at 24 °C in the dark. Embryos were then transferred to WCI containing 10 mg L^−1^ hygromycin, and incubated at 24 °C in the dark, subsequently being subcultured onto fresh WCI with hygromycin selection at 10 mg L^−1^, every 2 weeks over the next 5 weeks. For the final (5^th^) week on WCI, cultures were maintained in low light conditions at 24 °C. Cultures were then transferred onto wheat regeneration medium (WRM), supplemented with 2.5 mg L^−1^ zeatin and 10 mg L^−1^ hygromycin, in deep petri dishes (90 mm diameter × 20 mm), and cultured under full fluorescent lights (100 μM m^−2^ s^−1^) with a 16 h photoperiod. Regenerated plantlets were transferred to De Wit culture tubes (Duchefa-Biochemie, W1607), containing rooting medium supplemented with 20 mg L^−1^ hygromycin. After approximately 10 days, rooted plants were transferred to soil (cereal mix in 24CT trays) and acclimatized as in Hayta et al., 2019. The transgenic plants were maintained under the same growing conditions as donor material, with a long-day photoperiod (16 h at 600 μmol m^−2^ s^−1^ light, at 20 °C day and 16 °C night). Transgenesis was confirmed and transgene copy number analysis performed using Taqman qPCR and probe as described in Hayta et al., (2019). Values obtained were used to calculate transgene copy number, according to published methods (Livak and Schmittgen, 2001).

### Screening for gene edits

Six T_0_ lines were chosen for analysis, and 12 plants per T_1_ line were grown and analysed. A further 32 plants (T_2_) from one T_1_ edited plant were analysed for the presence of gene edits. Two sets of primers were designed to amplify the two target regions within the *TaDMC1-D1* CDS. For amplicon 1 (TaDMC1-D Guide1 target area), primers were TaDMC1-D F1 5′-GAGCGTGGGCTTGGTGTTAC-3′ and TaDMC1-D R1 5′-GAGGCGGAAGCACCCGGG-3′. For Amplicon 2 (TaDMC1-D Guide2 target area), TaDMC1-D F2 5′-TGCGATAGAATCTTCTGAAGTTTGTGTA-3′ and TaDMC1-D R2 5′-TCAATCCCTCCTTCAAATTACGC-3′ were used. PCR amplification was performed using GoTaq® Master Mix (Promega, M7122), with the following conditions: 3 min 94 °C, 40 cycles of 30 s at 94 °C, 15 s at 58 °C, 1 min at 72 °C and 5 min at 72 °C. Amplicons were Sanger sequenced directly (using their respective forward primers) by the Molecular Genetics Platform at the John Innes Centre.

### KASP genotyping of *ttmei1* mutants

Wild-type Chinese Spring and *ttmei1* mutant plants were grown to the 2-3 leaf stage, and DNA extracted from leaf material, as in Draeger et al. 2020 (adapted from Pallotta et al., 2003). Final DNA template concentrations were between 15-30 ng. KASP genotyping was performed using 5D chromosome-specific KASP primers with homeologous SNPs at the 3′ end, previously selected from the Wheat Breeders’ 820 K Axiom® array (Winfield et al., 2015), available at www.cerealsdb.uk.net, and aligned with the Chinese Spring reference sequence assembly, IWGSC RefSeq v1.0, (International Wheat Genome Sequencing Consortium, 2018). Two KASP primers were used to identify the *ttmei1* deletion region: BA00822801, based on a marker mapping proximal to *DMC1-D1*, and BA00750321, mapping distal to *DMC1-D1*. Primer sequences are shown in Table 1. The allele-specific forward primers and common reverse primers were synthesized by Merck https://www.merckgroup.com/. Allele-specific primers were synthesized with standard FAM or VIC compatible tails at their 5’ ends (see Table 1).

**Table 1.**
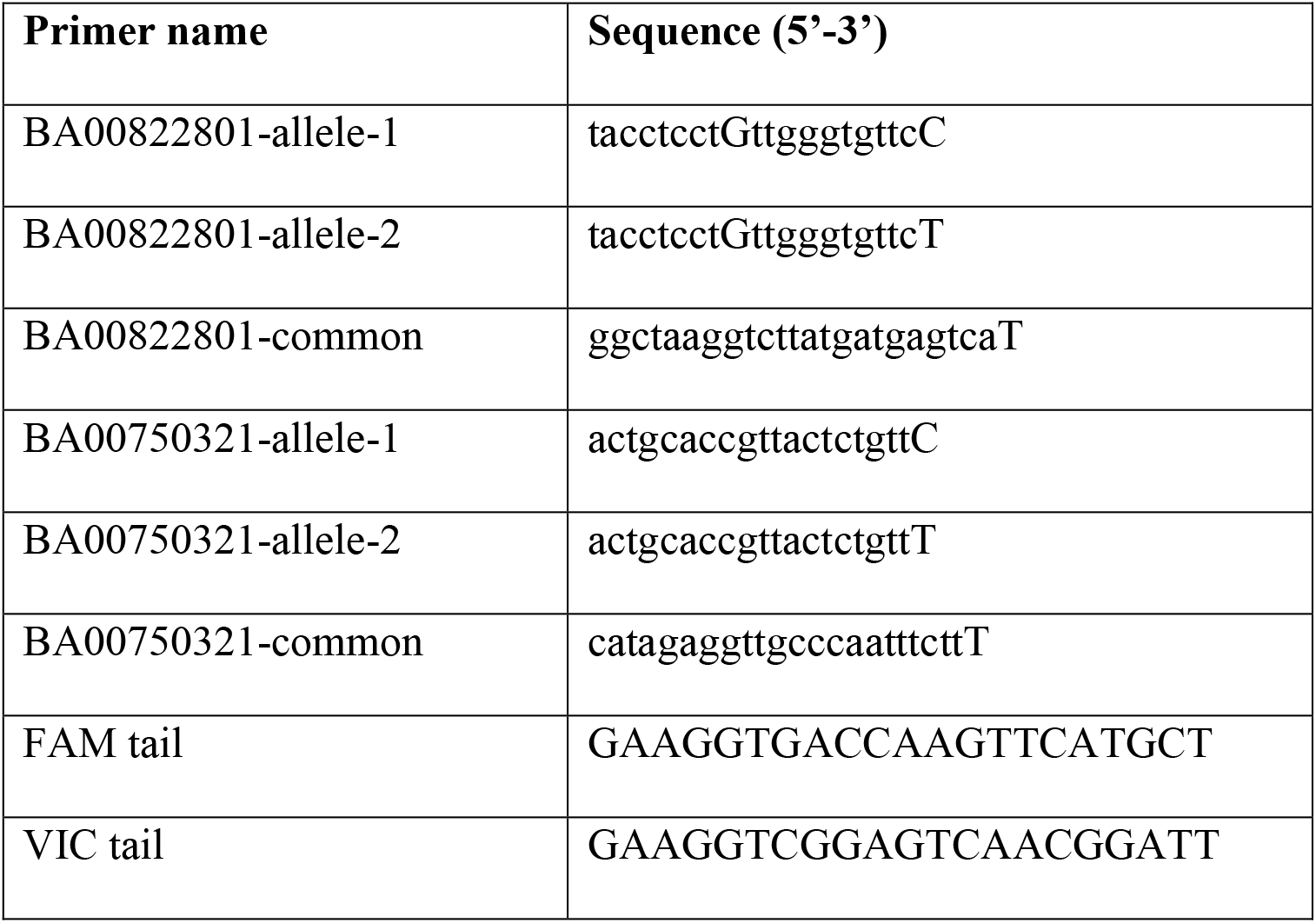
Primers for KASP genotyping of *ttmei1* (*Tadmc1-D1*) mutants.

### KASP reaction and PCR conditions

The KASP reaction and its components were as recommended by LGC Genomics Ltd and described at https://www.biosearchtech.com/support/education/kasp-genotyping-reagents/how-does-kasp-work.

Assays were set up as 5 μl reactions in a 384-well format, and included 2.5 μl genomic DNA template (15-30 ng of DNA), 2.5 μl of KASP 2x Master Mix (LGC Genomics) and 0.07 μl primer mix. Primer mix consisted of 12 μl of each tailed primer (100 μM), 30 μl common primer (100 μM) and 46 μl dH_2_O. PCR amplification was performed using the following programme: Hotstart at 94 °C for 15 min, followed by ten touchdown cycles (94 °C for 20 s; touchdown from 65-57 °C for 1 min, decreasing by 0.8 °C per cycle), followed by 30 cycles of amplification (94 °C for 20 s; 57 °C for 1 min). Fluorescent signals from PCR products were read in a PHERAstar microplate reader (BMG LABTECH Ltd.). If tight genotyping clusters were not obtained, additional rounds of amplification were performed. Genotyping data was analysed using KlusterCaller software (LGC Genomics).

### Analysis of *dmc1-D1* and *ttmei1* mutants at meiotic metaphase I

CRISPR *dmc1-D1* and *ttmei1* (Bc_2_F_2_) mutants and Chinese Spring control plants were initially grown in pots in a controlled environment room at 20°C day and 15°C night, with a 16-hour photoperiod and 70 % humidity, until development of the main shoot or tiller to be sampled had progressed to Zadoks growth stage 39 (Zadoks et al., 1974; Tottman, 1987), when the flag leaf ligule was just visible and meiocytes were deemed to be at premeiotic interphase. At this stage, the immature spikes enclosed within the leaf sheaths were between 3.5-6.5 cm in length (average 4.8 cm). Plants were then transferred to growth cabinets under continuous light and exposed to a low temperature (13 °C) for 6-7 days (with 70% humidity) or a high temperature (30 °C) for 24 h (75% humidity). For the high temperature experiments, treatments were initiated at a similar time of day, between 11.00 and 11.30 am, with the plant pots placed in trays of water to prevent dehydration.

Immediately following treatment, to identify anthers with metaphase I meiocytes, one anther from each floret was stained with acetocarmine and squashed to extrude the meiocytes, which were then examined using a DM2000 light microscope (Leica Microsystems). As the three anthers within a floret are synchronized in meiotic development, when metaphase I chromosomes were identified in one anther, the two remaining anthers from the same floret were prepared for cytological analysis by Feulgen staining with Schiff’s reagent, as described by Draeger et al., (2020). Anthers were sampled from three plants of each genotype, and images of metaphase I chromosomes captured using a DM2000 microscope equipped with a DFC450 camera and controlled by LAS v4.4 system software (Leica Microsystems). Images were captured in up to 8 different focal planes to aid scoring. For each plant, a minimum of 30 meiocytes were blind scored from digital images. This involved counting the following different meiotic chromosome configurations in each meiocyte: unpaired univalents (0 chiasmata), rod bivalents (1-2 chiasma), ring bivalents (2-3 chiasmata), trivalents (2–3 chiasmata), tetravalents (3 chiasmata) and pentavalents (4 chiasmata). Chiasma frequency per meiocyte was calculated separately using two different methods, with single chiasmata scores representing the minimum number of chiasmata per cell and double chiasmata scores representing the maximum. Figure 1 shows examples of the scored structures.

**Figure 1.**
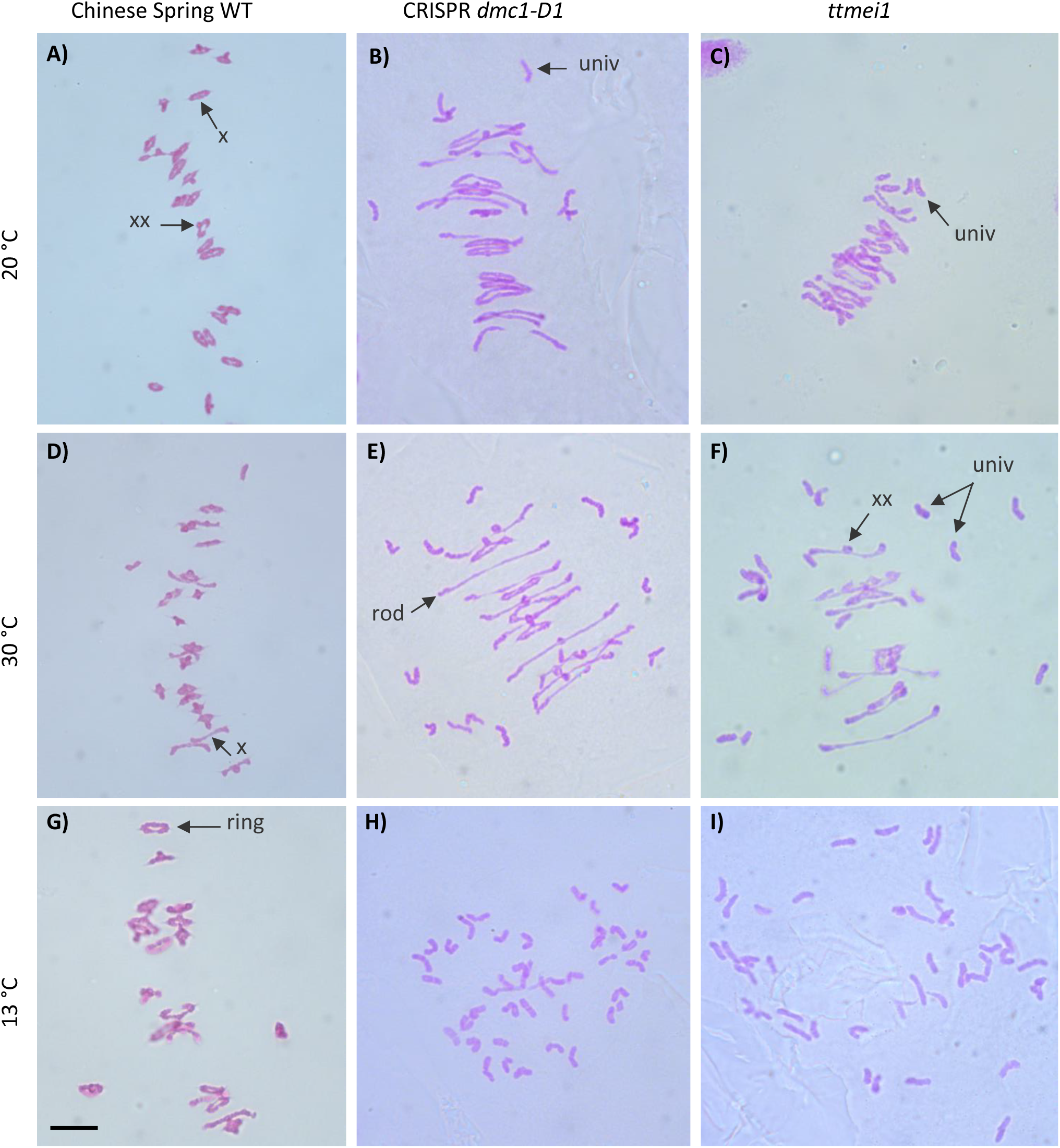
Representative images of Feulgen-stained metaphase I chromosomes from meiocytes of wild-type Chinese Spring, CRISPR *dmc1-D1* and *ttmei1* mutant plants treated at different temperatures. A) wild type, B) CRISPR *dmc1-D1* and C) *ttmei1* at normal temperatures; D) wild type, E) CRISPR *dmc1-D1* and F) *ttmei1* after 24 h at 30 °C; G) wild type, H) CRISPR *dmc1-D1* and I) *ttmei1* after 6-7 days at 13 °C. Examples of univalent chromosomes (univ), rod bivalents (rod), ring bivalents (ring), single chiasma (X) and double chiasmata (XX) are indicated with arrows; note complete univalence in CRISPR *dmc1-D1* and *ttmei1* mutants after treatment at 13 °C. Scale bars, 10 μm.

Statistical analyses were performed using STATISTIX 10.0 software (Analytical Software, Tallahassee, FL, USA). All treatments were analysed by the Kruskal–Wallis test (nonparametric one-way analysis of variance). Means were separated using Dunn’s test with a probability level of 0.05. Statistical analysis was carried out between genotypes (Table 2), and between temperatures (Table 3). Column charts were plotted using Microsoft Excel (2016) (Figure 2 and Figure 3).

**Table 2.** Effects of genotype on meiotic metaphase I chromosomes of wild-type Chinese Spring, CRISPR *Tadmc1-D1* and *ttmei1* mutant plants after treatment at 20 °C, 13 °C and 30 °C. The mean numbers of univalents, rod and ring bivalents, trivalents, tetravalents and pentavalents per meiocyte were scored along with single and double chiasma frequency. Standard error (SE) values are shown. P values < 0.05 indicate significant differences; lower case letters a-c indicate where the significant differences lie. For scores with the same letter, the difference between the means is not statistically significant. If the scores have different letters, they are significantly different. Ranges (in brackets) represent the minimum and maximum number of chromosomes with a particular configuration.

| Genotype | Temp.<br>(°C) | Univalent<br>Mean ± SE<br>(range) | Bivalent<br>(Rod)<br>Mean ± SE<br>(range) | Bivalent<br>(Ring)<br>Mean ± SE<br>(range) | Trivalent<br>Mean ± SE<br>(range) | Tetravalent<br>Mean ± SE<br>(range) | Pentavalent<br>Mean ± SE<br>(range) | Single<br>chiasmata<br>Mean ± SE<br>(range) | Double<br>chiasmata<br>Mean ± SE<br>(range) |
| --- | --- | --- | --- | --- | --- | --- | --- | --- | --- |
| 20°C day,<br>15°C<br>night | CS<br>wild type | 0.17 ± 0.12b<br>(0-2) | 1.18 ± 0.19b<br>(0-4) | 19.73 ± 0.21a<br>(16-21) | 0.00 ± 0.00<br>(0-0) | 0.00 ± 0.00<br>(0-0) | 0.00 ± 0.00<br>(0-0) | 40.66 ± 0.28a<br>(36-42) | 43.65 ± 0.32a<br>(38-47) |
|  | CRISPR<br>*dmc1-D1* | 4.25 ± 0.34a<br>(0-20) | 6.22 ± 0.24a<br>(1-15) | 12.66 ± 0.35b<br>(3-20) | 0.00 ± 0.00<br>(0-0) | 0.00 ± 0.00<br>(0-0) | 0.00 ± 0.00<br>(0-0) | 31.53 ± 0.49b<br>(14-41) | 34.16 ± 0.50b<br>(15-44) |
|  | *ttmei1*<br>Bc<sub>2</sub>F<sub>2</sub> | 3.89 ± 0.29a<br>(0-12) | 6.66 ± 0.19a<br>(1-12) | 12.38 ± 0.25b<br>(5-18) | 0.02 ± 0.01<br>(0-1) | 0.00 ± 0.00<br>(0-0) | 0.00 ± 0.00<br>(0-0) | 31.44 ± 0.36b<br>(22-39) | 33.24 ± 0.40b<br>(24-41) |
| p-value |  | < 0.0001 | < 0.0001 | < 0.0001 | - | - | - | < 0.0001 | < 0.0001 |
| 13°C,<br>6-7 days | CS<br>wild type | 0.24 ± 0.10b<br>(0-4) | 1.16 ± 0.25c<br>(0-7) | 19.74 ± 0.26a<br>(13-21) | 0.00 ± 0.00<br>(0-0) | 0.00 ± 0.00<br>(0-0) | 0.00 ± 0.00<br>(0-0) | 40.65 ± 0.28a<br>(33-42) | 43.62 ± 0.38a<br>(37-48) |
|  | CRISPR<br>*dmc1-D1* | 38.82 ± 0.19a<br>(34-42) | 1.57 ± 0.09b<br>(0-4) | 0.02 ± 0.01b<br>(0-1) | 0.00 ± 0.00<br>(0-0) | 0.00 ± 0.00<br>(0-0) | 0.00 ± 0.00<br>(0-0) | 1.61 ± 0.10b<br>(0-4) | 1.82 ± 0.11b<br>(0-6) |
|  | *ttmei1*<br>Bc<sub>2</sub>F<sub>2</sub> | 37.37 ± 0.32a<br>(24-42) | 2.19 ± 0.15a<br>(0-8) | 0.12 ± 0.03b<br>(0-2) | 0.00 ± 0.00<br>(0-0) | 0.00 ± 0.00<br>(0-0) | 0.00 ± 0.00<br>(0-0) | 2.44 ± 0.17b<br>(0-10) | 2.59 ± 0.19b<br>(0-11) |
| p-value |  | < 0.0001 | < 0.0001 | < 0.0001 | - | - | - | < 0.0001 | < 0.0001 |
| 30°C,<br>24 hours | CS<br>wild type | 0.21 ± 0.11c<br>(0-4) | 2.00 ± 0.25b<br>(0-5) | 18.90 ± 0.26a<br>(16-21) | 0.00 ± 0.00<br>(0-0) | 0.00 ± 0.00<br>(0-0) | 0.00 ± 0.00<br>(0-0) | 39.79 ± 0.29a<br>(35-42) | 42.44 ± 0.34a<br>(37-46) |
|  | CRISPR<br>*dmc1-D1* | 10.86 ± 0.52b<br>(0-26) | 8.12 ± 0.20a<br>(2-13) | 7.43 ± 0.27b<br>(1-17) | 0.02 ± 0.01b<br>(0-1) | 0.00 ± 0.00<br>(0-0) | 0.00 ± 0.00<br>(0-0) | 23.01 ± 0.49b<br>(10-37) | 25.50 ± 0.52b<br>(11-40) |
|  | *ttmei1*<br>Bc<sub>2</sub>F<sub>2</sub> | 20.67 ± 0.47a<br>(4-36) | 7.77 ± 0.17a<br>(3-16) | 2.75 ± 0.15c<br>(0-12) | 0.07 ± 0.02a<br>(0-2) | 0.01 ± 0.01<br>(0-1) | 0.01 ± 0.01<br>(0-1) | 13.47 ± 0.35c<br>(3-29) | 14.54 ± 0.37c<br>(3-32) |
| p-value |  | < 0.0001 | < 0.0001 | < 0.0001 | 0.0057 | - | - | < 0.0001 | < 0.0001 |

**Table 3.** Effects of three different temperature treatments on meiotic metaphase I chromosomes of wild-type Chinese Spring, CRISPR *Tadmc1-D1* and *ttmei1* mutant plants. The mean numbers of univalents, rod and ring bivalents, trivalents, tetravalents and pentavalents per meiocyte were scored along with single and double chiasma frequency. Standard error (SE) values are shown. P values < 0.05 indicate significant differences; lower case letters a-c indicate where the significant differences lie. For scores with the same letter, the difference between the means is not statistically significant. If the scores have different letters, they are significantly different. Ranges (in brackets) represent the minimum and maximum number of chromosomes with a particular configuration.

| Genotype | Temp.<br>(°C) | Univalent<br>Mean ± SE<br>(range) | Bivalent<br>(Rod)<br>Mean ± SE<br>(range) | Bivalent<br>(Ring)<br>Mean ± SE<br>(range) | Trivalent<br>Mean ± SE<br>(range) | Tetravalent<br>Mean ± SE<br>(range) | Pentavalent<br>Mean ± SE<br>(range) | Single<br>chiasmata<br>Mean ± SE<br>(range) | Double<br>chiasmata<br>Mean ± SE<br>(range) |
| --- | --- | --- | --- | --- | --- | --- | --- | --- | --- |
| CS<br>wild type | 20°C day,<br>15°C night | 0.17 ± 0.12<br>(0-2) | 1.18 ± 0.19b<br>(0-4) | 19.73 ± 0.21a<br>(16-21) | 0.00 ± 0.00<br>(0-0) | 0.00 ± 0.00<br>(0-0) | 0.00 ± 0.00<br>(0-0) | 40.66 ± 0.24a<br>(36-42) | 43.65 ± 0.32a<br>(38-47) |
|  | 13°C,<br>6-7 days | 0.24 ± 0.10<br>(0-4) | 1.16 ± 0.25b<br>(0-7) | 19.74 ± 0.26a<br>(13-21) | 0.00 ± 0.00<br>(0-0) | 0.00 ± 0.00<br>(0-0) | 0.00 ± 0.00<br>(0-0) | 40.65 ± 0.28a<br>(33-42) | 43.62 ± 0.38a<br>(37-48) |
|  | 30°C,<br>24 hours | 0.21 ± 0.11<br>(0-4) | 2.00 ± 0.25a<br>(0-5) | 18.90 ± 0.26b<br>(16-21) | 0.00 ± 0.00<br>(0-0) | 0.00 ± 0.00<br>(0-0) | 0.00 ± 0.00<br>(0-0) | 39.79 ± 0.29b<br>(35-42) | 42.44 ± 0.34b<br>(37-46) |
| p-value |  | **0.9964** | **< 0.0001** | **< 0.0001** | - | - | - | **< 0.0001** | **< 0.0001** |
| CRISPR<br>*dmc1-D1* | 20°C day,<br>15°C night | 4.25 ± 0.34c<br>(0-20) | 6.22 ± 0.24b<br>(1-15) | 12.66 ± 0.35a<br>(3-20) | 0.00 ± 0.00<br>(0-0) | 0.00 ± 0.00<br>(0-0) | 0.00 ± 0.00<br>(0-0) | 31.53 ± 0.49a<br>(14-41) | 34.16 ± 0.50a<br>(15-44) |
|  | 13°C,<br>6-7 days | 38.82 ± 0.19a<br>(34-42) | 1.57 ± 0.09c<br>(0-4) | 0.02 ± 0.01c<br>(0-1) | 0.00 ± 0.00<br>(0-0) | 0.00 ± 0.00<br>(0-0) | 0.00 ± 0.00<br>(0-0) | 1.61 ± 0.10c<br>(0-4) | 1.82 ± 0.11c<br>(0-6) |
|  | 30°C,<br>24 hours | 10.86 ± 0.52b<br>(0-26) | 8.12 ± 0.20a<br>(2-13) | 7.43 ± 0.27b<br>(1-17) | 0.02 ± 0.01<br>(0-1) | 0.00 ± 0.00<br>(0-0) | 0.00 ± 0.00<br>(0-0) | 23.01 ± 0.49b<br>(10-37) | 25.50 ± 0.52b<br>(11-40) |
| p-value |  | **< 0.0001** | **< 0.0001** | **< 0.0001** | - | - | - | **< 0.0001** | **< 0.0001** |
| *ttmei1*<br>Bc<sub>2</sub>F<sub>2</sub> | 20°C day,<br>15°C night | 3.89 ± 0.29c<br>(0-12) | 6.66 ± 0.19b<br>(1-12) | 12.38 ± 0.25a<br>(5-18) | 0.02 ± 0.01<br>(0-1) | 0.00 ± 0.00<br>(0-0) | 0.00 ± 0.00<br>(0-0) | 31.44 ± 0.36a<br>(22-39) | 33.24 ± 0.40a<br>(24-41) |
|  | 13°C,<br>6-7 days | 37.37 ± 0.32a<br>(24-42) | 2.19 ± 0.15c<br>(0-8) | 0.12 ± 0.03c<br>(0-2) | 0.00 ± 0.00<br>(0-0) | 0.00 ± 0.00<br>(0-0) | 0.00 ± 0.00<br>(0-0) | 2.44 ± 0.17c<br>(0-10) | 2.59 ± 0.19c<br>(0-11) |
|  | 30°C,<br>24 hours | 20.67 ± 0.47b<br>(4-36) | 7.77 ± 0.17a<br>(3-16) | 2.75 ± 0.15b<br>(0-12) | 0.07 ± 0.02<br>(0-2) | 0.01 ± 0.01<br>(0-1) | 0.01 ± 0.01<br>(0-1) | 13.47 ± 0.35b<br>(3-29) | 14.54 ± 0.37b<br>(3-32) |
| p-value |  | **< 0.0001** | **< 0.0001** | **< 0.0001** | - | - | - | **< 0.0001** | **< 0.0001** |

**Figure 2.**
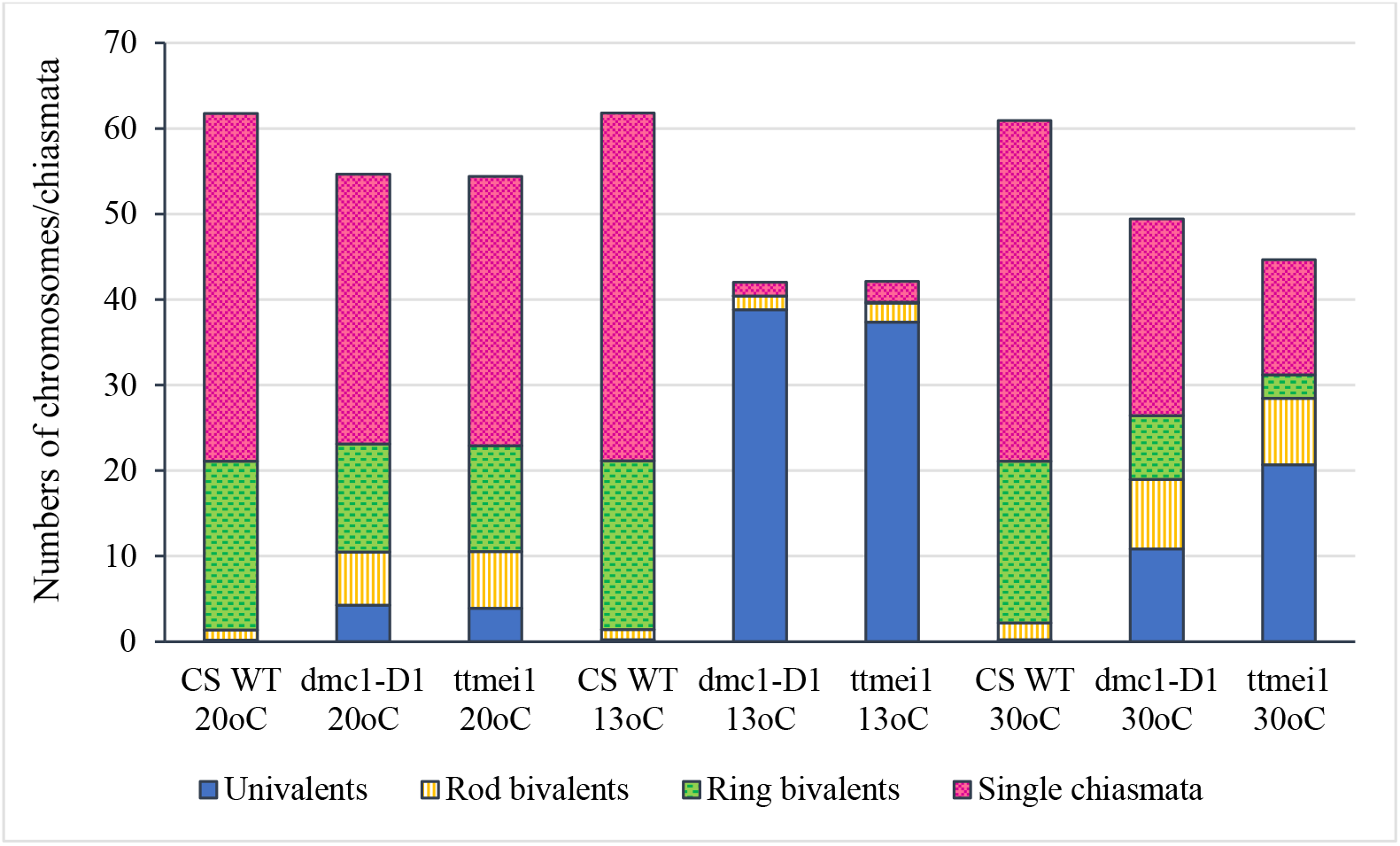
Column chart showing the effects of genotype on meiotic metaphase I chromosomes of CRISPR *dmc1-D1* and *ttmei1* mutants compared with Chinese Spring wild type (CS WT). Numbers of univalents, rod and ring bivalents and single chiasmata are shown. Numbers of multivalents and double chiasmata are not shown. Note reduced chiasma frequencies in *dmc1-D1* and *ttmei1* mutants, particularly at 13°C.

**Figure 3.**
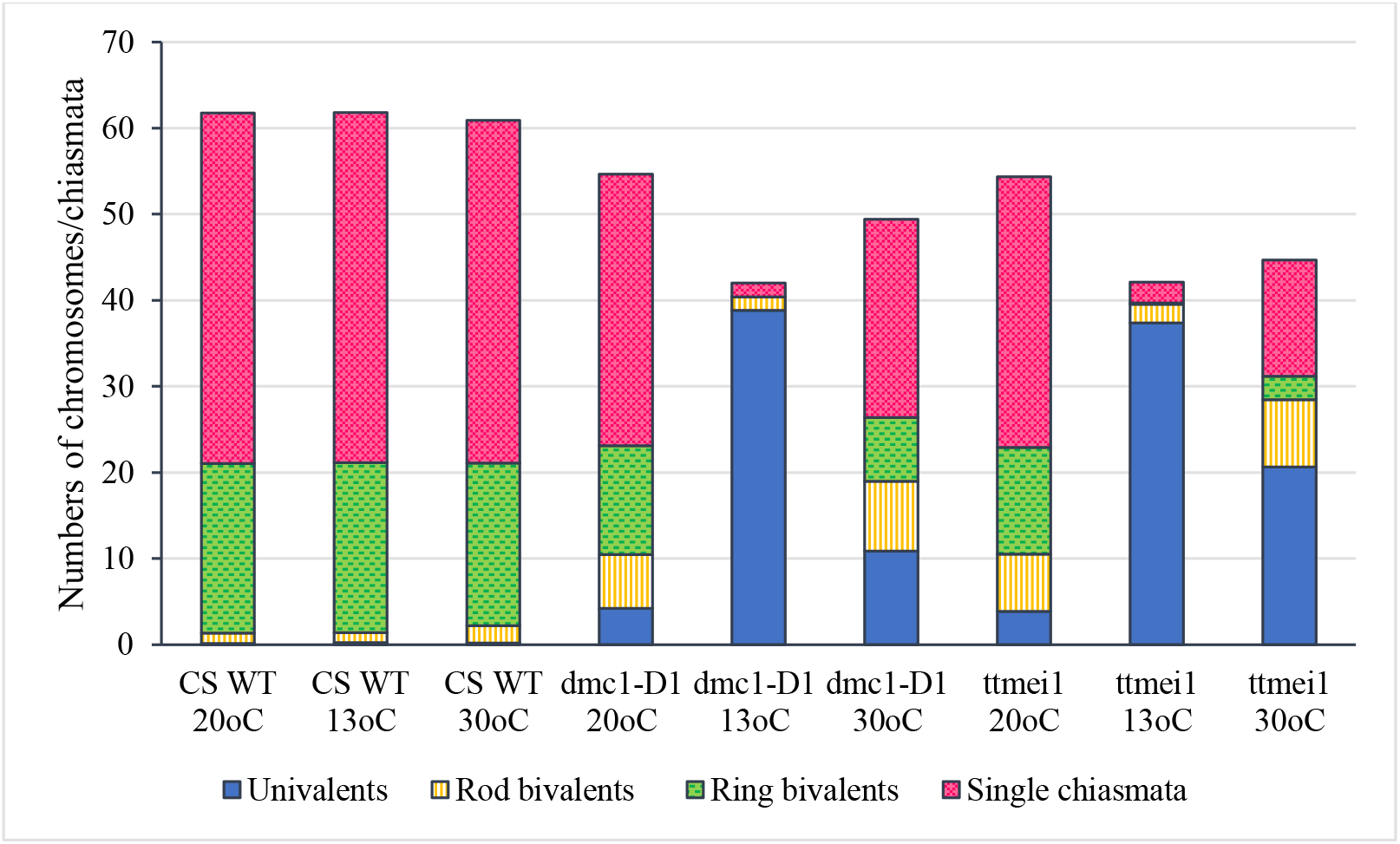
Column chart showing the effects of three different temperature treatments (20 °C, 13 °C and 30 °C) on meiotic metaphase I chromosomes of Chinese Spring wild type (CS WT) plants and CRISPR *dmc1-D1* and *ttmei1* mutants. Numbers of univalents, rod and ring bivalents and single chiasmata are shown. Numbers of multivalents and double chiasmata are not shown. Note reduced chiasma frequencies in *dmc1-D1* and *ttmei1* mutants at 30°C and almost total univalence at 13°C.

## Results

### CRISPR *Tadmc1-D1* mutants

From each of 6 selected T_0_ lines, 12 T_1_ lines were analysed for edits in the *TaDMC1-D1* target region. Sanger sequencing revealed that none of these lines had a homozygous edit, but one T_1_ plant had a heterozygous edit in the target region. The edited T_1_ plant was self-fertilized, and 32 T_2_ progeny plants were sequenced. In 9 of the T_2_ generation plants, sequencing revealed a 39 bp homozygous deletion within the *DMC1-D1* target region. Examples of the T_2_ sequence chromatograms are shown in Supplementary Figure 1. These 9 T_2_ plants were used in the temperature treatment experiments: 3 plants were treated at 13 °C for 6-7 days; 3 were treated at 30 °C for 24 h; and 3 control plants remained under normal control conditions of 20 °C day, 15 °C night.

### Reduction in chiasma frequency in *dmc1-D1* and *ttmei1* mutants at normal temperatures

Meiotic metaphase I chromosomes were blind scored in 3 plants of each genotype at each of 13 °C, 30 °C and normal temperatures. A minimum of 30 meiocytes were scored for each plant (at least 100 meiocytes per genotype). At normal temperatures, wild-type plants contained ∼20 ring bivalents and a single rod bivalent per meiocyte as usual, with univalents occurring only occasionally (Figure 1A; Table 2; Figure 2). The mean number of chiasmata was 41-44. However, under the same temperature conditions, in the *dmc1-D1* and *ttmei1* mutants, univalents (∼4) and rod bivalents (6-7) occurred significantly more frequently than in the wild-type plants, whereas ring bivalents (12-13), single chiasmata (31-32) and double chiasmata (33-34) were significantly fewer than in the wild type (Figure 1B and C; Table 2; Figure 2). No multivalent chromosomes were observed at normal temperatures in either wild-type or *dmc1-D1* meiocytes. In one *ttmei1* mutant plant, two trivalents were observed, but this was not significantly different to the wild-type or *dmc1-D1* scores.

### Further reduction in chiasma frequency at 30 °C in *dmc1-D1* and *ttmei1* mutants

After 24 hours at 30 °C, in wild type plants, mean numbers of univalents per meiocyte remained the same as at normal temperatures (< 1), whereas numbers of rod bivalents increased from around one to around two, ring bivalents decreased from ∼20 to ∼19, and single and double chiasmata were correspondingly reduced, which were all significant differences, albeit small (Figure 1A and D; Table 3; Figure 3). In the *dmc1-D1* mutants, numbers of univalents and rod bivalents increased from ∼4 and ∼6 respectively at normal temperatures to ∼11 and ∼8 respectively after treatment at 30 °C; numbers of ring bivalents decreased from ∼13 to ∼7, single chiasmata numbers decreased from ∼32 to ∼23 (a reduction of ∼27%) and double chiasmata from ∼34 to ∼26 (a reduction of ∼25%), which were all significant differences (Figure 1B and E; Table 3; Figure 3). In *ttmei1* mutants, numbers of univalents and rod bivalents also increased at 30 °C, from ∼4 to ∼21 and from ∼7 to ∼8 respectively; ring bivalents decreased from 13 to ∼3, single chiasmata dropped from ∼31 to ∼14 (a reduction of ∼57%) and double chiasmata from ∼33 to ∼15 (a reduction of ∼56%) (Figure 1C and F; Table 3; Figure 3). Again, all differences were significant. Although numbers of rod bivalents were similar in both mutants at 30 °C, differences between numbers of univalents, ring bivalents and chiasmata at normal temperatures and at 30 °C were larger in *ttmei1* mutants than in *dmc1-D1* mutants. At 30 °C, a small but significant number of trivalents were observed in *dmc1-D1* and *ttmei1* mutants, but not in wild-type plants. Most of the trivalents were observed in a single *ttmei1* plant. A single tetravalent and two pentavalents were observed in the same *ttmei1* mutant, but this was not a significant difference. No tetravalents or pentavalents were observed in any other plants.

### Crossover failure at 13 °C in *dmc-D1* and *ttmei1* mutants

After 6-7 days treatment at 13 °C, in wild-type plants, numbers of univalents, rod and ring bivalents and chiasmata per meiocyte were similar those seen at normal temperatures (Figure 1A and C; Table 3; Figure 3). However, in the *dmc1-D1* mutants, after treatment at 13 °C, there was a dramatic increase in the number of univalents from ∼4 to ∼39; ring bivalent numbers decreased from ∼13 to almost none (< 1), rod bivalents decreased from ∼6 to ∼2 and chiasma frequency decreased dramatically from ∼32 (single chiasmata) and ∼34 (double chiasmata) to only ∼2. Almost all observed chromosomes (∼39 out of 42) were univalents (Figure 1H; Table 3; Figure 3). Scores for *ttmei1* mutants were similar to those for *dmc1-D1* mutants: 37 out of 42 chromosomes were univalents, there were almost no ring bivalents (< 1), around two rod bivalents and an average of only 2-3 chiasmata per meiocyte (Figure 1I; Table 3; Figure 3). This means that, at 13 ° C, crossover was reduced by ∼96% in *dmc1-D1* mutants and by ∼94% in *ttmei1* mutants, compared to levels seen in wild-type plants at the same temperature. Furthermore, in *dmc1-D1* mutants, exposing plants to 13 °C for 6-7 days reduced crossover by ∼95% compared to levels observed at normal temperatures. In *ttmei1* mutants, the reduction in crossover was 92%. No multivalent chromosomes were observed in any plants at 13°C.

## Discussion

### Development of CRISPR *dmc1* mutants in wheat

Using RNA-guided Cas9, we have developed new CRISPR mutants containing a 39 bp deletion in the 5D copy of the *DMC1* gene in the hexaploid wheat reference variety Chinese Spring. Until recently, wheat transformation has remained genotype dependent, therefore limiting the potential use of genomic tools such as CRISPR-Cas technologies. However, recent development and deployment of the morphological gene fusion GRF-GIF (Debernardi et al., 2020), coupled with our efficient and robust transformation system (Hayta et al., 2019, 2021), has reduced genotype dependence in wheat and enabled us to report this first CRISPR-Cas9 targeted mutagenesis in Chinese Spring. These CRISPR *dmc1-D1* mutants, along with backcrossed *ttmei1* mutants (containing a 4 Mb deletion of *DMC1-D1)*, were used to determine whether meiosis is stabilized by *DMC1-D1* at high and/or low temperatures.

### Reduction in crossover in *dmc1-D1* mutants at normal temperatures

In wild type plants grown at normal temperatures, chromosomes aligned on the equatorial plate as normal, pairing as bivalents, mostly rings, but with the occasional rod bivalent (Figure 1A). However, in the *ttmei1* and CRISPR *dmc1-D1* mutants there were significantly more univalents and rod bivalents, and significantly fewer ring bivalents and chiasmata (Figure 1B and C; Table 2; Figure 2), with a reduction in chiasma (and therefore crossover) frequency of 22-24%. Two multivalents were observed in one *ttmei1* mutant, but none in the other *ttmei1* mutants or in any of the *dmc1-D1* mutants, so the deletion of *DMC1-D1* does not seem linked to the occurrence of multivalents. Clearly, disruption of *DMC1-D1* has a significant effect on meiosis, but this effect is not severe at normal temperatures, probably due to gene redundancy, since previous studies have shown that homeologs of the *DMC1* gene on chromosomes 5A and 5B are present and expressed (Draeger et al., 2020).

### Crossover is reduced by 95% in *dmc1-D1* mutants at 13 °C, and synapsis does not complete

In the current study, Chinese Spring wild-type plants and *dmc1-D1* and *ttmei1* mutants were exposed to a low temperature treatment of 13 °C for 6-7 days during a period lasting from premeiotic interphase to early meiosis I. Exposure to the low temperature had no significant effect on metaphase I chromosomes in the wild type plants (Table 3, Figure 3), but in the CRISPR *dmc1-D1* mutants, almost all chromosomes observed were univalents (Figure 1G and H), and crossover decreased by over 95% compared to that seen in the wild-type plants (Table 2, Figure 2). Similar results were obtained for the *ttmei1* mutants (Figure 1I). In the *dmc1-D1* mutants, exposure to the low temperature reduced crossover by ∼95% compared to that observed at normal temperatures, and in the *ttmei1* mutants crossover was reduced by 92%. The low chiasma frequencies and high numbers of univalent chromosomes observed in the *dmc1-D1* and *ttmei1* mutants at low temperatures were similar to those observed when the whole of chromosome 5D is deleted in Chinese Spring (Draeger et al., 2020).

This high number of univalent chromosomes suggests a major problem with crossover formation. Consistent with this, previously we showed that *ttmei1* mutants exhibit significant abnormalities of synapsis at low temperatures (Draeger et al., 2020): after exposure to a low temperature of 13 °C, in both wild type and *ttmei1* mutant meiocytes, synapsis initiates normally at one pole of the nucleus at early zygotene, but in the mutant, synapsis does not complete at pachytene. These experiments confirm our previous hypothesis that, in wheat, *DMC1-D1* is responsible for the preservation of normal synapsis and crossover at lower temperatures, and is therefore equivalent to the *Ltp1* locus first described by Hayter and Riley in 1967, providing an answer to a question that has existed for over 55 years.

Most reported SC failures involve high temperatures, but, as in wheat, low-temperature failures have also been reported in *Hyacinthus orientalis* and two species of *Solanum* (Elliott, 1955; Karihaloo, 1991). Moreover, in the ectothermic Japanese red-bellied newt, *Cynops pyrrhogaster*, low temperatures during meiosis also give rise to univalent chromosomes, indicating failure of chromosome pairing due to asynapsis, and leading to abnormal spermatozoa production (Yazawa et al., 2003). *DMC1* expression in *C. pyrrhogaster* also decreases under the same low-temperature conditions (8 °C or 12 °C), suggesting its low level of expression may contribute to the temperature-dependent abnormalities seen in spermatogenesis. This study supports our suggestion that *DMC1* is involved in the maintenance of normal chromosome synapsis at low temperatures.

### Reduction in crossover in *dmc1-D1* mutants at high temperatures

In the current study, exposure of *dmc1-D1* mutants to a high temperature of 30 °C for just 24 hours during premeiotic interphase to meiosis I, resulted in a reduced number of crossovers and increased univalence, though to a lesser extent than that seen after a low temperature treatment of 13 °C. Similar results were obtained for *ttmei1* mutants, although there was more variation between scores for individual plants. Previously, high variation between chiasma frequency scores was also observed in the original *ttmei1* mutant plants (prior to backcrossing), following treatment at 30 °C for 24 hours (Draeger et al., 2020). It was suggested that this variation could be linked to a high level of background mutations due to gamma irradiation. In the current study, backcrossing these mutants with Chinese Spring for two generations (Bc_2_F_2_), should have substantially reduced the large numbers of undesirable background mutations, but variation between scores was still high, suggesting that this variation is less likely to be due to background mutations.

High variation between scores for different plants may have occurred if the heat applied reached the meiocytes at slightly different developmental stages or for longer or shorter durations. When Chinese Spring plants are grown at 20 °C under continuous light, meiosis is estimated to take around 24 hours to complete (Bennett et al., 1971, 1973), though at higher (25 °C) temperatures, meiosis is accelerated and completes in around 18 hours (Bennett et al., 1972). Although assigning meiosis to specific growth stages by assessing the external morphology of a plant is unreliable (Barber et al., 2015), in the current study, the 24-hour high temperature treatments should have been of sufficient duration to coincide with the temperature-sensitive period from premeiotic interphase to early meiosis I (as described in Bayliss and Riley, 1972b, and Draeger and Moore, 2017). Other studies have suggested that it might be difficult to deliver a heat stress treatment to cells such as meiocytes at a specific time, because within an anther they are surrounded by many different cell layers, such as the tapetum layer and the epidermis, which are able to buffer the high temperatures. However, in *Arabidopsis thaliana*, tracking the deposition of stress granules which form at elevated temperatures has demonstrated that the ambient temperature reaches the meiocytes in less than 15 minutes (De Jaeger-Braet et al., 2022). Even so, since the structures in wheat are much larger, it is still possible that a change in temperature may take longer to reach different meiocytes, due to their varying level of insulation according to the location of their anther within a spike.

### The role of *DMC1* in meiosis

*TaDMC1-D1* expression is at its highest during early meiotic prophase I (Draeger et al., 2020), which in wheat, is when synapsis initiates at the ‘telomere bouquet’ stage (Martín et al., 2017). DMC1 has a central role in synapsis and homologous recombination. It is a meiosis-specific protein, structurally similar to the bacterial strand-exchange recombinase, RECA (Bishop et al., 1992). Homologous recombination is initiated by programmed DNA DSBs at leptotene, which results in single-stranded DNA ‘overhangs’ at the break sites. DMC1 and RAD51 (another RECA homolog) form helical nucleoprotein filaments by polymerizing on the single-stranded overhangs. These filaments perform homology searches and carry out strand invasion and strand exchange between homologous chromosomes (Neale and Keeney, 2006). Repair of these interhomolog invasion events results in crossovers or non-crossovers, although only a small minority of DSBs are repaired as crossovers (Reviewed in Lambing et al., 2017). RAD51 has a role in both somatic and meiotic repair, and is essential for maintaining chromosomal integrity in mitotic cells. DMC1 is the main catalytically active strand-exchange protein during meiosis, but it is also thought to suppress RAD51-mediated recombination in plant meiosis (Da Ines et al., 2022).

*DMC1* homologs are found in a wide variety of organisms. In yeast (*Saccharomyces* cerevisiae) and mice, *dmc1* deficiency results in defective meiotic recombination and chromosome synapsis, with cells arresting in prophase, leading to sterility (Bishop et al., 1992; Pittman et al., 1998; Yoshida et al., 1998; Bannister et al., 2007). Similarly, in wheat *Tadmc1-D1* (*ttmei1*) mutants, synapsis does not complete, and meiosis appears to arrest before pachytene in late prophase, although this was after a treatment at 13 °C (Draeger et al., 2020), whereas the phenotypes observed in yeast and mice were at ambient temperatures. Disruption of *DMC1* also leads to sterility in most diploid plant species. *Arabidopsis thaliana* has a single copy of the *DMC1* gene, and synapsis is disrupted in *Atdmc1* mutants, which also show high levels of univalence, and drastically reduced fertility (Couteau et al., 1999). Rice (*Oryza sativa*) has two *DMC1* homologs, *OsDMC1A* and *OsDMC1B* (Ding et al., 2001). A mutation in either one of these homologs does not cause problems during meiosis, but synapsis is abnormal in *Osdmc1a Osdmc1b* double mutants, and they also exhibit serious crossover defects, high numbers of univalents at metaphase and are sterile (Wang et al., 2016). Barley (*Hordeum vulgare*) has a single *DMC1* homolog, *HvDMC1*, and mutations in this gene lead to abnormal synapsis, multiple univalents and chromosome mis-segregation (Colas et al., 2019; Szurman-Zubrzycka et al., 2019). As in yeast and mice, disruption of the barley orthologue of *DMC1* at ambient temperatures leads to a phenotype similar to that of *Tadmc1* at low temperatures.

### The contribution of *DMC1* homeologs to temperature tolerance

From our own results, it appears that, in Chinese Spring, *TaDMC1*-*D1* promotes low temperature tolerance, and possibly high temperature tolerance. There are two other homeologs of the *DMC1* gene in hexaploid wheat: *TaDMC1-A1* (TraesCS5A02G133000) on chromosome 5A and *TaDMC1-B1* (TraesCS5B02G131900) on 5B. All three copies are expressed in wheat (Devisetty et al., 2010). Previously, we found differences in gene expression levels between the three homeologs in meiotic anthers, with *DMC1-D1* having the highest meiotic gene expression levels and *DMC1-A1* the lowest (Draeger et al., 2020). It is not yet known how the 5A and 5B copies contribute to meiosis, but these differences in gene expression could be related to differences in the abilities of these three genes to stabilize the genome at low temperatures.

Tetraploid wheat (AABB) has only two copies of *DMC1* (Tang et al., 2017), but synapsis at 12 °C is normal, despite the absence of chromosome 5D (Riley et al., 1966). This is probably due to the presence of a dominant *Ltp* allele on chromosome 5A, with a similar chromosome stabilizing activity to that of chromosome 5D in hexaploid wheat (Hayter and Riley, 1967). Interestingly, some other varieties and subspecies of wheat differ from Chinese Spring, in that the gene responsible for stabilizing chromosome pairing at low temperatures is located on chromosome 5A rather than 5D (Chapman and Miller, 1981). Future research will require development of *dmc1-A1* and *dmc1-B1* mutants, along with double and triple mutants in different hexaploid and tetraploid wheat varieties, to determine how each of the *DMC1* homeologs contributes to stabilizing synapsis and crossover at high and low temperatures. This could be achieved using CRISPR/Cas9, which, in addition to its high specificity, can be used to simultaneously target multiple copies of a gene, a technology that has already enabled the production of loss-of-function triple wheat mutants (Taagen et al, 2020; Li et al, 2021). Recently, another genome editing system, transcription activator-like effector nucleases (TALENs), has been used to disrupt all six *CmDMC1* loci in the hexaploid flower, *Chrysanthemum morifolium* (Shinoyama et al., 2020).

### Dosage effects of *TaDMC1* alleles

Different dosages of the *TaDMC1* alleles can affect the stability of synapsis and crossover at low temperatures. In Chinese Spring plants lacking *DMC1-D1*, the chromosome 5A and 5B homologs *DMC1-A1* and *DMC1-B1*, are unable to compensate for the lack of *DMC1-D1*, and cannot stabilize synapsis and crossover at low temperatures. In a previous study, even when chromosome 5B was present as a double dose in Chinese Spring plants, it was still unable to compensate for the lack of 5D, but when chromosome 5A was present as a double dose, chromosome pairing at 12 °C was normal (Riley et al., 1966). Since *TaDMC1*-*A1* has the lowest meiotic gene expression of the three *DMC1* homeologs in Chinese Spring, this suggests that when *DMC1*-*A1* is present as a double dose, an increase in its expression compensates for the loss of *DMC1-D1* to preserve low temperature tolerance.

Climate change is likely to have a negative effect on meiosis, and therefore on fertility and crop production, so screening of germplasm collections to identify heat-tolerant genotypes is a high priority for future crop improvement. It will also be important to determine the relative meiotic temperature tolerance of plants carrying these specific *TaDMC1* alleles growing under natural conditions.

## Author contribution statement

TD grew and maintained the plants, made the crosses, carried out the KASP genotyping, sampled anthers and collected metaphase I images for scoring the phenotype, produced the corresponding figure and wrote the manuscript. M-DR scored chromosome crossover, performed the statistical analysis and produced the graphs. SH and MS developed the CRISPR *Tadmc1-D1* mutant in ‘Chinese Spring’ using RNA-guided Cas9 and produced the sequence chromatograms figure; GM provided the concept, and with AM, provided thoughts and guidance, and revised and edited the manuscript.

## Acknowledgements

We thank the following John Innes Centre staff members: Martha Clarke of the Crop Transformation facility for her excellent technical support in the development of the CRISPR *dmc1-D1* mutant; Saleha Bakht, Molecular Genetics Platform Manager for sequencing services; the Germplasm Resource Unit for providing seed and the Horticultural Services Department for plant maintenance. We also acknowledge The International Atomic Energy Agency, Vienna for gamma irradiation of Chinese Spring seed to produce the original *ttmei1* mutant.

