## Supplementary Figure 1 for "*DMC1* stabilizes synapsis and crossover at high and low temperatures during wheat meiosis"

### Slide 1
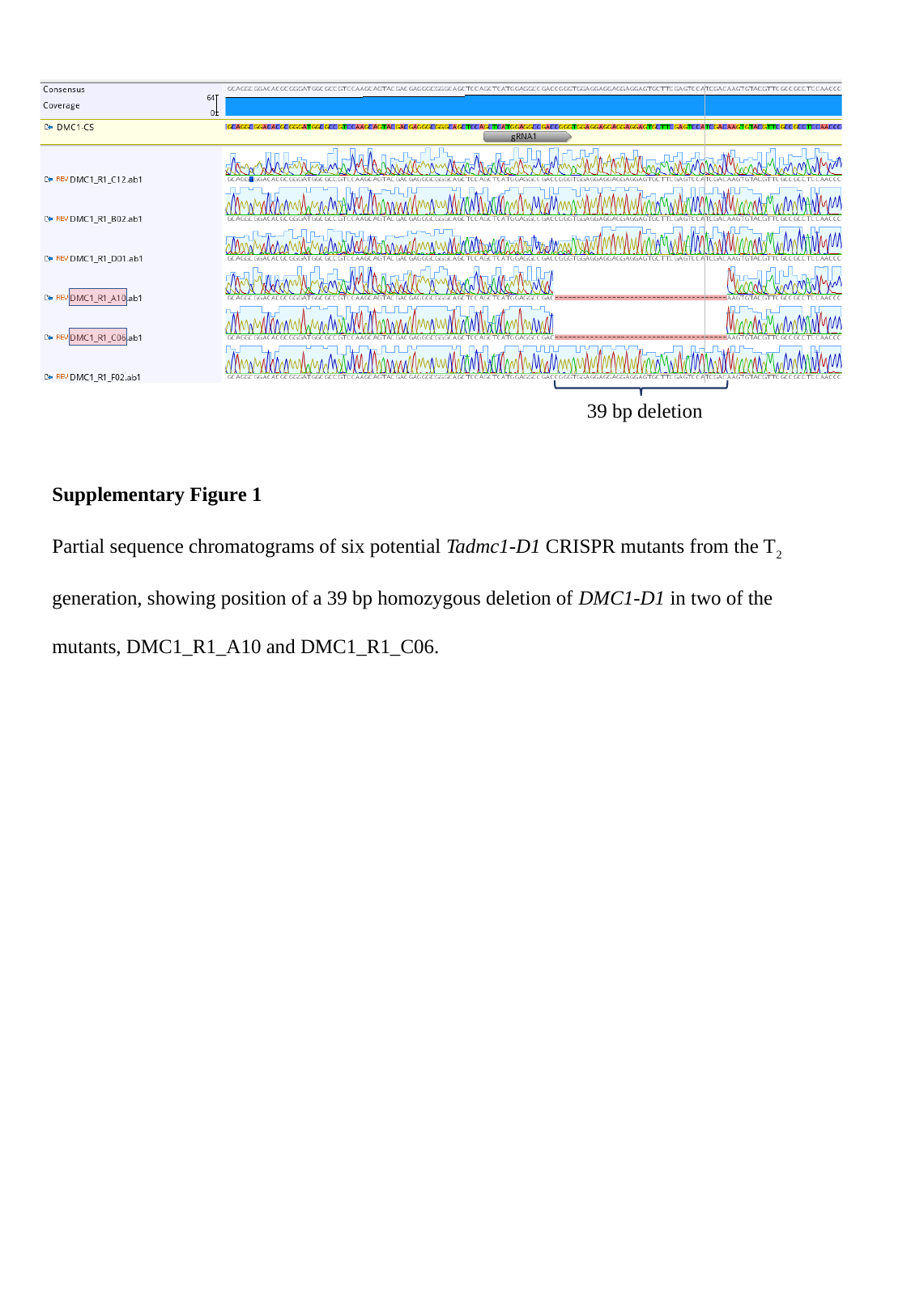

39 bp deletion
Supplementary Figure 1
Partial sequence chromatograms of six potential Tadmc1-D1 CRISPR mutants from the T_2_ generation, showing position of a 39 bp homozygous deletion of DMC1-D1 in two of the mutants, DMC1_R1_A10 and DMC1_R1_C06.
